# Complement C3 recognition by C3 convertases

**DOI:** 10.1101/2025.08.09.669456

**Authors:** Changhao Jia, Xiaoke Yang, Ming-hui Zhao, Ying Tan, Junyu Xiao

**Author notes:** Correspondence: Ming-hui Zhao, Ying Tan, Junyu Xiao. These authors contributed equally to this work (Changhao Jia, Xiaoke Yang).

## Abstract

The complement system plays a fundamental role in human immunity, and its dysregulation is implicated in numerous diseases. Activation of the complement occurs through three main pathways: classical, lectin, and alternative; which converge at the central component, C3. The classical and lectin pathways utilize the C4b2a convertase to cleave C3 and initiate complement activation, while the alternative pathway employs the C3bBb convertase, which is further stabilized by properdin. The molecular mechanisms governing C3 recognition by these convertase complexes remain incompletely understood. Here, we first present a 3.1 Å cryo-electron microscopy (cryo-EM) structure of the C4b2a–C3 complex, alongside the 2.9 Å and 3.1 Å structures of the C4b2 zymogen in the loading and activation states. These structural snapshots elucidate the structural basis for C3 engagement by C4b2a, and illustrate sequential conformational changes during the classical/lectin pathway convertase maturation. Furthermore, we determine a 2.6 Å cryo-EM structure of the C3bBb–properdin–C3 complex, which uncovers unique substrate-binding features of C3bBb and sheds light on how properdin stabilizes the alternative pathway convertase. These results offer comprehensive mechanistic insights into complement activation.

## Main Text

The complement system is a vital component of innate immunity, comprising a complex network of over 50 soluble proteins and cell surface receptors that play essential roles in pathogen clearance, immune modulation, and the maintenance of homeostasis (*1, 2*). Dysregulation of the complement system has been associated with various human disorders, including renal, rheumatic, neurological, and cardiovascular diseases, etc. (*3–6*). More recently, complement dysregulation has also been implicated in SARS-CoV-2-related pathogenesis, particularly with respect to thrombosis (*7–8*).

The complement system operates through three distinct but interconnected activation pathways: the classical pathway, the lectin pathway, and the alternative pathway. Each of these pathways converges at the central component, C3, which is cleaved by the C3 convertases to produce the fragments C3a and C3b. C3a functions as an anaphylatoxin, triggering inflammatory responses through its interaction with the G protein-coupled receptor C3aR, while C3b acts as an opsonin that binds to pathogens, enhancing their recognition and clearance by phagocytes. Additionally, C3b serves as a critical subunit for the alternative pathway C3 convertase, promoting the amplification of the complement response.

C3 convertases are key enzymatic complexes within the complement system that play a crucial role in activating the complement cascade (Fig. 1A). There are two main forms of C3 convertase: C4b2a (here the convention where ’a’ denotes the enzymatically active fragment of C2 is used) in the classical and lectin pathways, and C3bBb in the alternative pathway (*9, 10*). The C4b2a convertase is formed through the binding of C4b, generated by the serine protease C1s in the classical pathway or MASP-2 in the lectin pathway, to C2, which is further cleaved by C1s or MASP-2, resulting in the active C4b2a form. In contrast, the C3bBb convertase arises from the binding of C3b to factor B (FB), which is activated by factor D (FD). The resulting C3bBb convertase is stabilized by properdin (P) (*11*). Alternatively, a small fraction of C3 can undergo spontaneous hydrolysis, known as "C3 tick-over", yielding C3(H2O) that adopts a C3b-like conformation (*12*), and can thereby lead to the formation of the C3(H2O)Bb convertase. Recent studies have shown that granzyme K (GZMK) can also activate the complement cascade by cleaving C4 and C2, which leads to the formation of the C4b2a convertase (*13*), or by directly cleaving and activating C3 (*14*).

**Fig. 1:**
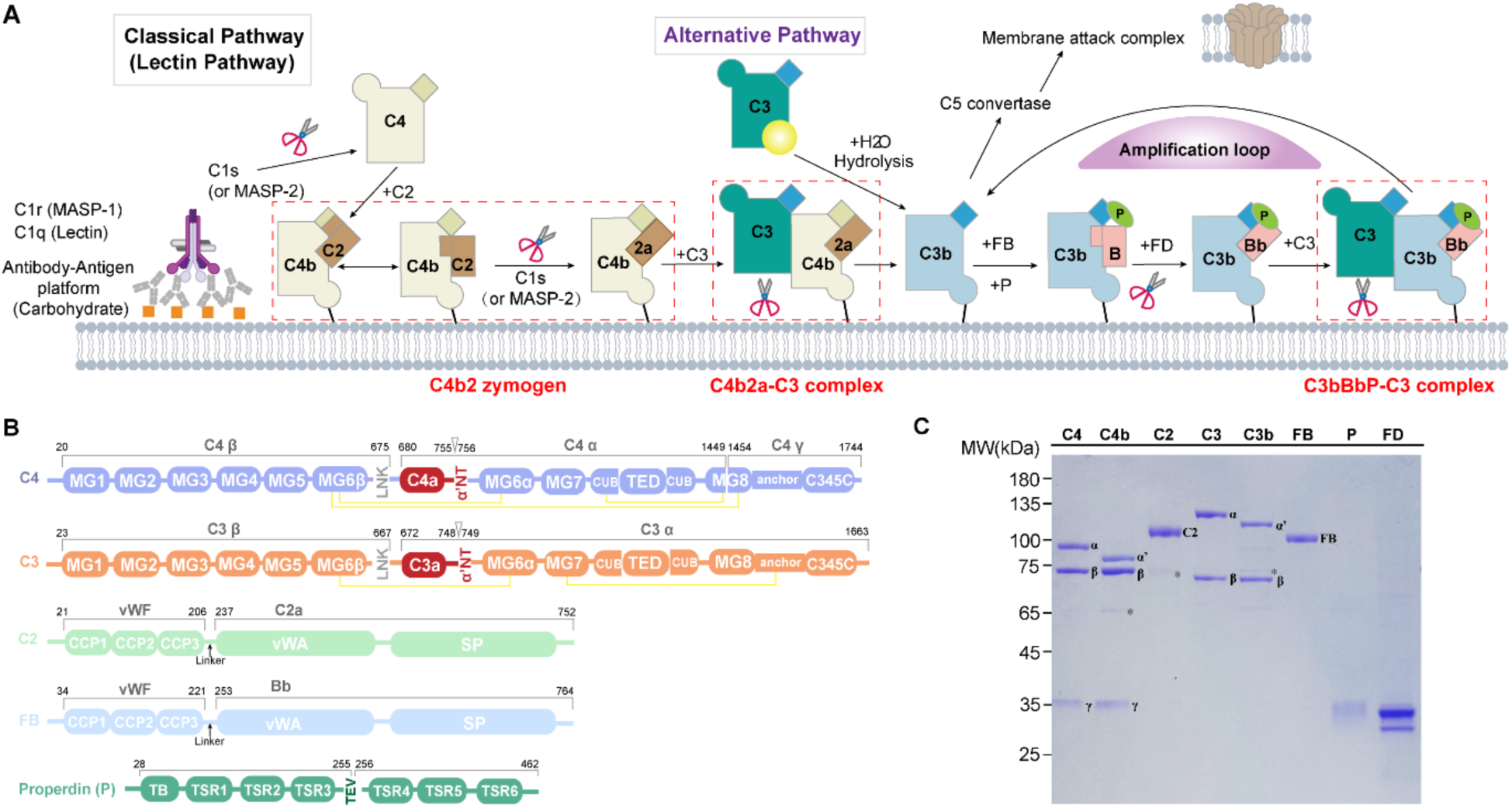
Overview of the complement C3 convertases. **(A)** A simplified diagram of the complement cascade. **(B)** Schematic representations of the domain organization of C3 convertase components, including C4, C2, C3, factor B (FB), and properdin. MG: Macroglobulin domain; LNK: Linker domain; α’-NT: C3 α’ chain N-terminal; CUB: Complement C1r-C1s, UEGF, BMP1; TED: Thioester domain; vWF: von Willebrand factor; vWA: von Willebrand A domain; SP: Serine protease domain. The yellow connecting lines indicate interchain disulfide bonds, while the triangular arrows denote cleavage sites on C4 and C3. **(C)** SDS-PAGE analysis of purified proteins, including C4, C4b, C2, C3, C3b, FB, properdin, and factor D (FD). Asterisks (*) indicate contaminant proteins.

Both C4b2a and C3bBb are inherently labile, dissociating within minutes after formation (*15, 16*), posing significant challenges for their study. Previous breakthroughs, in particular the determination of the C3bB zymogen structure in complex with factor D (*17*) and the C3bBb– SCIN (Staphylococcal complement inhibitor) complex (*18*), have provided foundational insights into the alternative pathway convertase assembly. The general location of the C3 substrate binding site on C3bBb was inferred from the crystallographic dimer interface formed between the two C3b molecules in the C3bBb–SCIN structure. However, mechanistic details underlying C3 recognition by the C3bBb convertase remain unclear. Similarly, the assembly and substrate engagement mechanisms of the classical/lectin pathway C4b2a convertase are poorly understood, limiting therapeutic targeting of these complexes.

In this study, we employ cryo-electron microscopy (cryo-EM) to resolve the molecular architecture of these convertases in functional states. We report a 3.1 Å structure of the C4b2a– C3 complex, elucidating the structural basis for C3 recognition by the classical/lectin pathway convertase. We also resolve the 2.9 Å and 3.1 Å structures of the C4b2 zymogen (standardized notation for the C4bC2 complex) in the loading and activation states, and thereby reveal sequential conformational transitions during the classical/lectin pathway convertase formation and maturation. Furthermore, we capture a 2.6 Å structure of the C3bBbP–C3 complex, which unveils distinct features governing substrate engagement by C3bBb. Detailed analysis also delineates how properdin stabilizes the C3b–Bb interaction, thereby positively regulating the alternative pathway convertase. These high-resolution snapshots address long-standing questions in complement biology, and provide a foundation for advancing the development of C3-targeted therapeutics.

## Cryo-EM structure of the C4b2a–C3 complex

To capture a stable C3 convertase for cryo-EM analysis, we first isolated C3 and C4 from human plasma as previously described (*19*) (Fig. 1B, C). C4b was then generated by incubating C4 with C1s. Additionally, we produced a catalytically inactive mutant of C2, containing the S679A mutation, using HEK293F cells. Following this, C4b, C2, and C3 were mixed in an equal molar ratio, with Ni^2+^ included to enhance the C4b–C2 interaction (*20*). Afterwards, this sample was briefly treated with C1s and then rapidly plunged in liquid ethane for cryo-EM analysis. The C1s treatment resulted in partial cleavage of C2, leading to the formation of the C4b2a convertase, which readily engages C3 and enables us to trap the classical pathway C3 convertase-substrate complex and visualize the structure at a resolution of 3.1 Å (Fig. 2A; Fig. S1; Table S1). Additionally, we captured two functional states of the C4b2 pro-convertase containing intact C2: a loading state at 2.9 Å and an activation state at 3.1 Å.

**Fig. 2:**
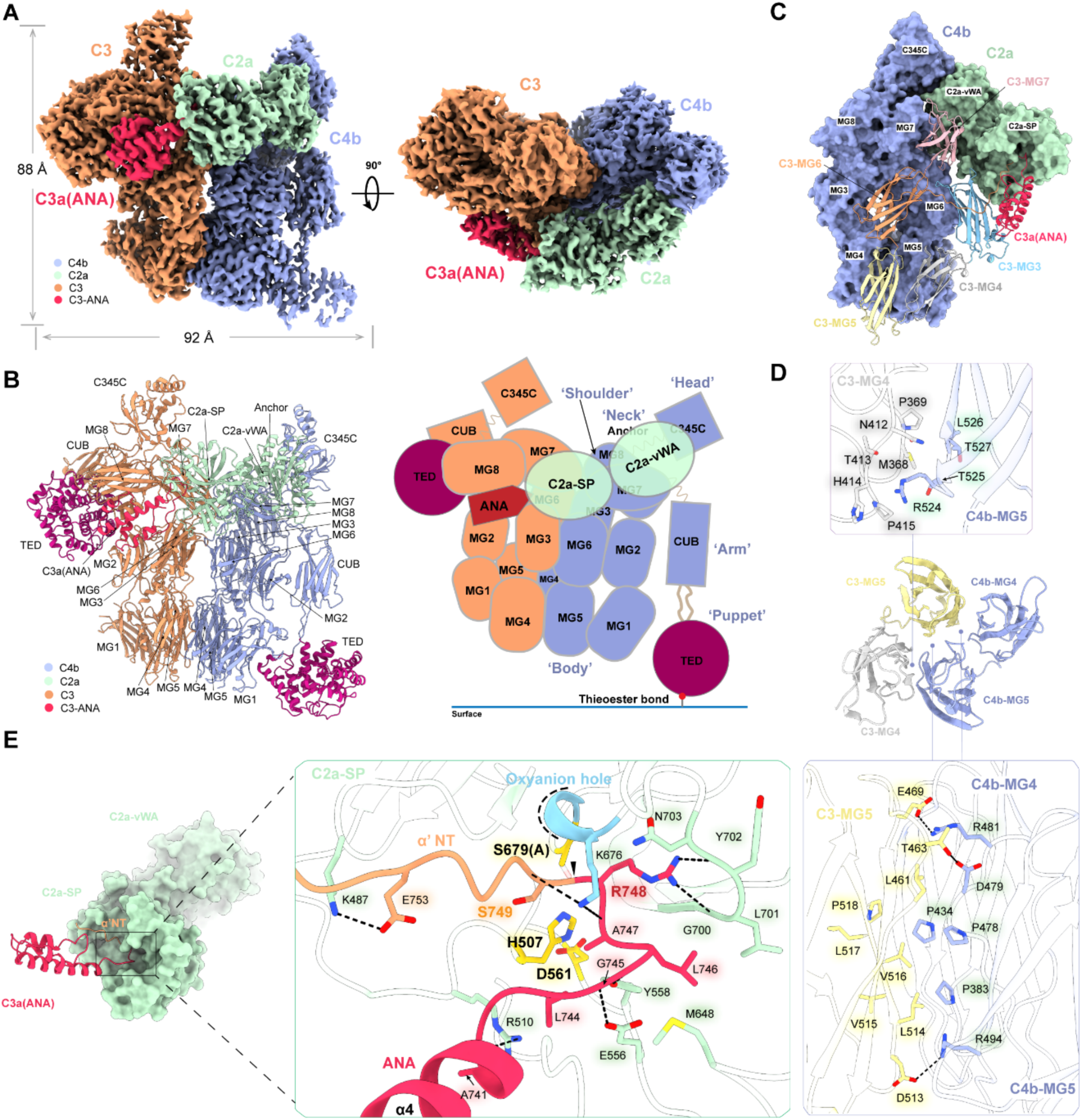
Cryo-EM structure of the classical/lectin pathway C3 convertase. **(A)** Two views of the cryo-EM reconstruction of the C4b2a–C3 complex. Complement proteins C4b, C3, and C2b are depicted in blue, orange, and green, respectively, while the C3a(ANA) domain is highlighted in red. **(B)** A ribbon diagram of the C4b2a–C3 structure accompanied by a schematic representation. **(C)** The interaction between C4b and C3 involves multiple MG domains from both proteins. **(D)** Interaction details between C3-MG4, C3-MG5 and C4b-MG5, C4b-MG4. **(E)** Interaction between the C3 scissile loop and the C2a-SP domain. C3a(ANA) and α’-NT are shown in red and orange, respectively. The cleavage site is indicated by a black arrowhead. The oxyanion hole in C2a-SP is highlighted in blue, and the catalytic triad Ser679(A), His507, Asp561 are highlighted in yellow. The hydroxyl group of Ser679 is hypothetical and is represented in a lighter color.

In the C4b2a–C3 complex, C4b adopts an active conformation typical of C3b and C4b, resembling a "puppeteer" structure (*21*) (Fig. 2B). This includes a head (C345C domain), a neck (anchor), shoulders (MG7–8), a body (MG1–6, LNK), and a downward-extending arm (CUB domain), with a "puppet" (TED domain) dropped to the level of MG1–MG4–MG5. Compared to the structure of C4b alone (*22*), the C345C domain exhibits a slight rotation to accommodate C2a (Fig. S2A). C2a binds to the C345C domain of C4b via its vWA domain, while its serine protease (SP) domain swings toward C3. The substrate C3 adopts a characteristic inactive conformation, with the TED domain securely tucked beneath the CUB and MG8 domains, effectively concealing the Cys1010–Gln1013 thioester warhead (*23*) (Fig. S2B). The ANA (anaphylatoxin) or C3a domain, consisting of four helices, points towards the SP domain of C2a, facilitating the docking of the α’-NT segment, which contains the Arg748– Ser749 cleavage site, into the catalytic center of C2a.

## Substrate recognition mechanism of C4b2a

The structure of the C4b2a–C3 complex illustrates the specific recognition mechanism employed by the classical/lectin pathway C3 convertase. Both C4b and C2a are involved in C3 recognition, contributing to the stringent substrate specificity. The interface between C4b and C3 buries 2140 Å^2^ from each molecule and involves multiple MG domains from both proteins (Fig. 2C). Specifically, C4b-MG4 interacts with C3-MG5, facilitated by residues Pro383_C4b_, Pro434_C4b_, and the Leu514_C3_–Pro518_C3_ strand (Fig. 2D). Between C4b-MG5 and C3-MG5, Pro478_C4b_ and Asp479_C4b_ form interactions with Leu461_C3_ and Thr463_C3_, while Arg481_C4b_ and Arg494_C4b_ engage with Glu469_C3_ and Asp513_C3_, respectively. Additional interactions occur between C4b-MG5 and C3-MG4, where Arg524_C4b_ is surrounded by Asn412_C3_–Pro415_C3_, and Thr525_C4b_–Leu526_C4b_ pack against Met368_C3_–Pro369_C3_. The C4b-MG6 domain, which consists of two split subdomains (MG6α and MG6β), interacts with three domains in C3 (Fig. S2C): Asp576_C4b_ forms a salt bridge with Lys264_C3_ in C3-MG3; Glu585_C4b_ bonds with Arg855_C3_ in C3-MG7; and Arg828_C4b_ interacts with Asp572_C3_ in C3-MG6. Additionally, C4b- MG7 and C3-MG7 align closely, with Arg831_C4b_, Glu832_C4b_, and Leu859_C4b_ accommodating Phe920_C3_. Overall, two dozen hydrogen bonds and salt bridge contacts are formed between C4b and C3, ensuring their specific interaction.

The interaction between C2a and C3 buries 1360 Å^2^ from each molecule. The α’-NT segment (residues 743–756), which contains the scissile loop of C3, exhibits clear densities and is positioned within the active site of C2a-SP (Fig. 2E; Fig. S2D). Arg510_C2a_ contacts the C-terminal end of the α4 helix in the ANA/C3a domain, facilitating the positioning of Leu744_C3_ (P5 site). Residues Glu556_C2a_, Tyr558_C2a_, and Lys676_C2a_ orient Gly745_C3_–Leu746_C3_–Ala747_C3_ (P4–P2 residues) through hydrogen bond interactions with their main-chain groups, and Met648_C2a_ further packs on Leu746_C3_. Gly700_C2a_–Asn703_C2a_, located at the beginning of the long loop 2, accommodate Arg748_C3_ (P1). Additionally, Lys676_C2a_ interacts with the main-chain of Ser749_C3_ (P1’), while Lys487_C2a_ forms a salt bridge with Glu753_C3_ (P5’). Notably, in the absence of C3 binding, Lys676_C2a_ is drawn to Ser698_C2a_, causing its main-chain carbonyl group to form a hydrogen bond with Ser679_C2a_, which distorts the oxyanion hole (*24*). Upon C3 binding, Lys676_C2a_ interacts with Ala747_C3_ and Ser749_C3_, leading to the establishment of a functional oxyanion hole (Fig. S2E). In addition to engaging the scissile loop of C3, C2a also anchors to the C3-MG3 domain (Fig. 2C). A pocket formed by Lys485_C2a_, Leu517_C2a_, Arg519_C2a_, and Glu532_C2a_ accommodates Arg290_C3_, while Trp529_C2a_ packs with His333_C3_ (Fig. S2F). As a result of these interactions with both C2a and C4b, C3-MG3, along with the upper tip of C3-MG4, undergoes a ∼25° rotation within the MG1–6 ring (Fig. S2B). Collectively, these interactions ensure the precise positioning of the Arg748_C3_–Ser749_C3_ peptide bond, priming it for cleavage by the Asp561_C2a_–His507_C2a_–Ser679(Ala)_C2a_ catalytic triad.

## Formation of the classical/lectin pathway C3 convertase

We also captured the structure of the C4b2 pro-convertase at the loading and activation states. Together with the C4b2a structure in complex with the C3 substrate, these structures allow us to visualize the formation of the C3 convertase at discrete steps (Fig. 3A).

**Fig. 3:**
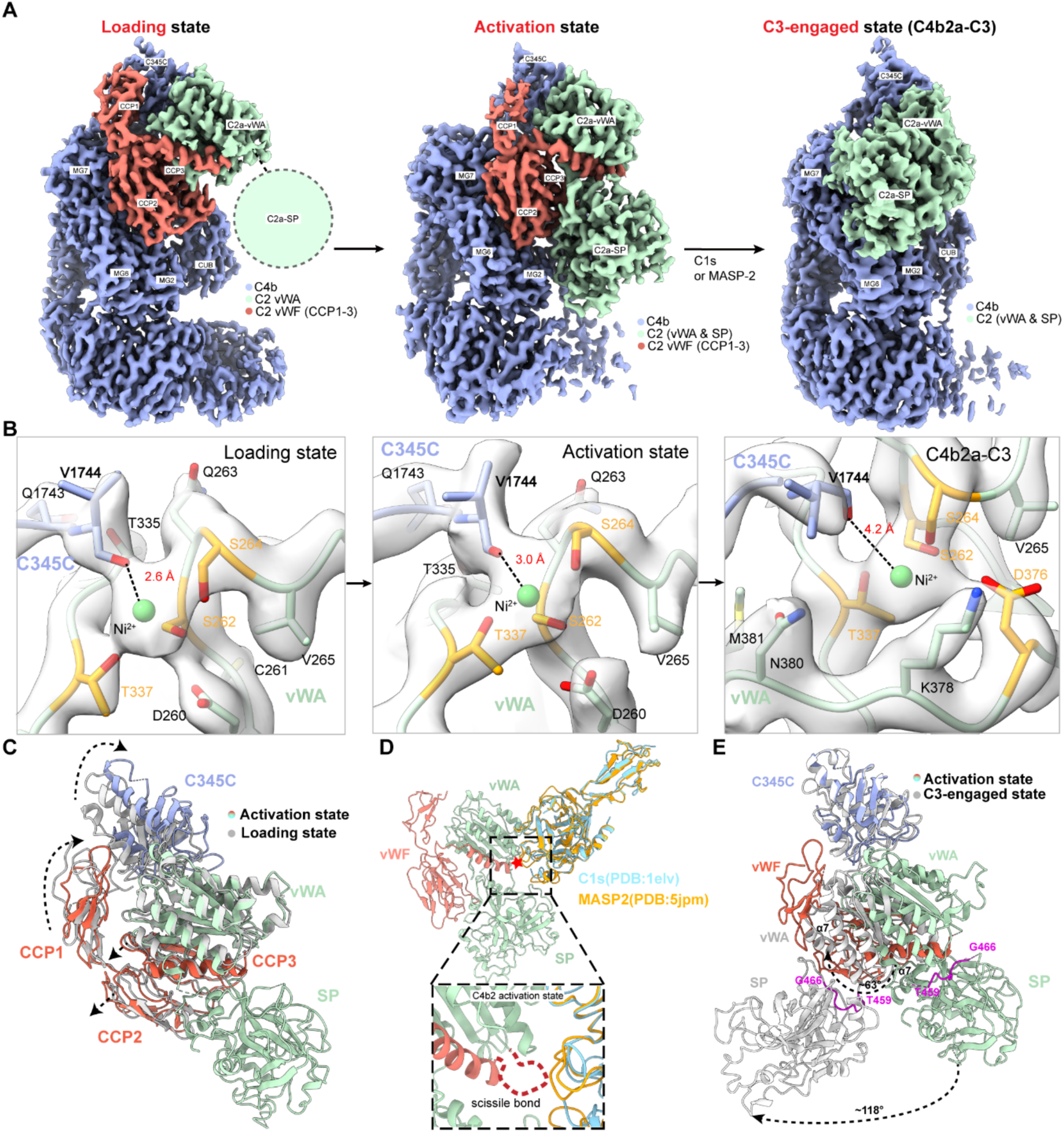
Formation of the classical/lectin pathway C3 convertase. **(A)** The C3 convertase at three distinct steps: loading, activation, and C3-engaged. C4b, C2b (CCP1–3), and C2a (vWA–SP) are depicted in purple, salmon and green, respectively. In the loading state, the C2a-SP domain exhibits flexibility and cannot be clearly modeled, and is represented by a dashed circle. **(B)** The positions of C4b-V1744 relative to the Ni^2+^ ion in C2-vWA. Ser262, Ser264, Thr337 and Asp376 in the MIDAS motif of C2-vWA are highlighted in orange. The density maps are shown at the same contour level. **(C)** Structural comparison between the loading and activation states of C4b2. The loading state is presented in gray, while the activation state uses the same color scheme as in panel a. **(D)** Structural models of C1s or MASP-2 bound to C2. These models are generated by aligning the activation state of the C4b2 structure with that of C3bBD (PDB: 2XWB) and then aligning the structures of C1s (PDB: 1ELV) and MASP-2 (PDB: 5JPM) to Factor D. **(E)** Structural comparison between the activation and C3-engaged states. The C3-engaged state is shown in gray, while the activation state is presented in color.

The loading state of the C4b2 structure exhibits similarities to the factor B complexed with cobra venom factor (CVF) (*25*), a homolog of C3b (Fig. S3A). In this state, the C2-vWF region, namely the CCP1–3 domains or the C2b pro-segment, and the C2-vWA domain interact with C4b. In particular, C2-CCP1 and C2-vWA together engage the C4b-C345C domain. Notably, akin to the interaction observed between CVF and FB, the carboxyl group of the terminal Val1744_C4b_ coordinates with the Ni^2+^ ion in the metal ion-dependent adhesion site (MIDAS) of C2-vWA, completing the metal ion coordination (Fig. 3B). Additionally, the CCP2–3 domains contact C4b-MG2, C4b-MG6, C4b-MG7, C4b-CUB, and the C4b-α’NT region (Fig. S3B). While the C2-SP domain exhibits flexibility in this state and cannot be clearly modeled despite the overall high resolution of the map, its general positioning suggests close proximity to the vWA domain (Fig. S3C). This arrangement would obstruct the access to the scissile site within the CCP3-vWA linker by the C1s protease, also reminiscent of the structural arrangement in the CVF-B complex.

The activation state of the C4b2 structure closely resembles the C3bB structure (*17*) (Fig. S3D), and provides a complete visualization of C2 (Fig. 3A). In this state, C2-SP adopts a stable conformation and is anchored to the MG2 and CUB domains of C4b. Additionally, the vWA-SP linker in C2 is more ordered, and is positioned between C2-vWA and C2-SP (Fig. S3E). This change pulls C2-vWA closer to C2-SP, which in turn induces a rotation of the C4b- C345C domain. As a result, the interaction between C2-CCP1 and C4b-C345C is reduced (Fig. 3C; Fig. S3F). This displacement likely contributes to the release of C2b pro-segment following C2 cleavage. Models of C1s or MASP2 can be generated based on Factor D, utilizing the C3bBD complex structure as a template (Fig. 3D). In the resulting models, the scissile loop would be readily accommodated in the active sites of these proteases, facilitating C2 cleavage and C4b2 convertase maturation.

In the C4b2a–C3 complex, C2a adopts a conformation closely resembling that of the C2a alone structure (*24*) (Fig. S3G), with the C2a-SP drawn towards the substrate C3 as described above. Consequently, a tectonic reorientation of C2a is observed when compared to the C2a region in the activation state of C4b2 (Fig. 3E, Movie 1). The C2a-vWA domain shifts into the position previously occupied by CCP1–3, while the C2a-SP undergoes a ∼118° swing. Thus, the CCP1-3 pro-segment must be released for the activated C2a to effectively dock onto C3. The large swing of the C2a-SP is further facilitated by a ∼63° rotation of the α7 helix in the C2a-vWA domain, accompanied by a complete reorientation of the vWA-SP linker (Fig. 3E, Movie 1). The C2a-vWA domain remains attached to the C4b-C345C domain. Notably, in the C3-bound state, the carboxyl group of Val1744_C4b_ is positioned further from the Ni^2+^ ion in the MIDAS motif of C2a-vWA, as evidenced by the density map (Fig. 3B). This suggests a loosened interaction between C4b and C2a, preparing C2a for release following C3 cleavage. It is likely that the smaller ionic radius of Mg^2+^ would further expedite this dissociation process *in vivo*. As such, engagement of C2a to C3 would prompt the concurrent release of C2a from C4b, explaining the labile nature of the C4b2a convertase.

## Cryo-EM structure of the C3bBbP–C3 complex

To investigate C3 recognition by the AP C3 convertase, we generated C3b from C3 through limited trypsin processing (*26*). A catalytically inactive factor B containing the S699A mutation was prepared using HEK293 cells. Additionally, we produced a two-chained monomeric properdin by inserting a Tobacco Etch Virus (TEV) protease cleavage site between its TSR3 and TSR4 domains (Fig. 1B), and then processing the purified properdin oligomer with TEV, as previously described (*27*). Using a similar approach to the generation of the C4b2a–C3 complex, we mixed purified C3b, factor B, properdin, and C3 in equal molar ratios with Ni^2+^. The mixture was then briefly treated with factor D, converting factor B into the active fragment Bb. Subsequently, cryo-EM analysis was conducted, resulting in the structure determination of the C3bBbP–C3 complex at an atomic resolution of 2.6 Å (Fig. 4A, B; Fig. S4; Table S1).

**Fig. 4:**
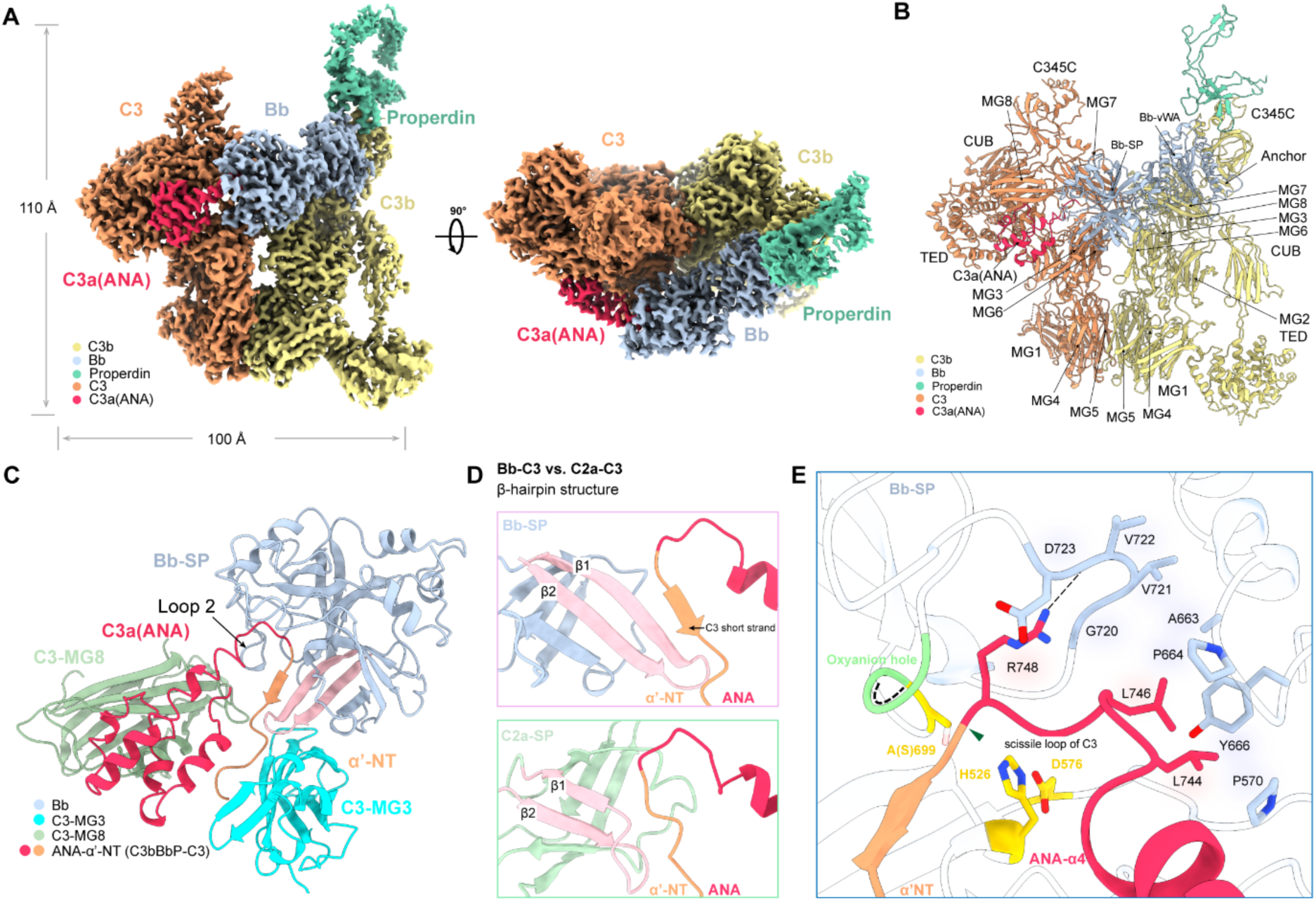
Cryo-EM structure of the alternative pathway C3 convertase. **(A)** Two views of the C3bBbP–C3 complex cryo-EM reconstruction. C3b, Bb, and C3 are depicted in khaki, light blue, and orange, respectively; while properdin is shown in aquamarine. C3a is highlighted in red. **(B)** Ribbon rendering of the C3bBbP–C3 structure. **(C)** Interaction between Bb-SP and C3, with Bb-SP, C3a, α’-NT, C3-MG3, C3-MG8 shown in light blue, red, orange, cyan, and dark green, respectively. The β-hairpin in Bb-SP is highlighted in pink. **(D)** The β-hairpin in Bb-SP is longer than that of C2a-SP, inducing the formation of a short strand in C3. **(E)** Interaction between the C3 scissile loop and Bb-SP.

In this complex, C3b and Bb individually closely resemble their respective structures observed in the C3bBb–SCIN (Staphylococcal complement inhibitor) complex (*18*) (Fig. S5A). It is apparent that Bb can now properly latch onto the substrate C3 without the steric hindrance imposed by SCIN. The C3 molecule adopts a conformation nearly identical to that seen in the C4b2a–C3 complex (Fig. S5B). Similar to the C4b2a–C3 complex, both C3b and Bb engage the C3 molecule. C3b forms a quasi-dimer with C3, involving the MG3–7 domains of both proteins. This C3b–C3 quasi-dimer shares similarities with the C3b–C3b homodimer observed in the C3bBb–SCIN crystal structure, especially in the lower region of the MG rings (Fig. S5C). However, the upper half of the MG ring including the MG2–3–6 domains in the C3 molecule, as well as the MG7 domain, undergoes rotations and leans further toward C3b in the C3b–C3 interaction. Compared to the C4b–C3 interface in the C4b2a–C3 structure, the C3b–C3 interaction involves fewer hydrogen bonds and salt bridge interactions (15 vs. 24), and the interface area is slightly smaller (2040 Å^2^ vs. 2140 Å^2^), suggesting that C4b forms a more optimized interaction with C3 than C3b.

In contrast, the interaction between Bb and C3 exhibits unique features and appears enhanced compared to the C2a–C3 interaction (Fig. 4C). For instance, beyond targeting the α’- NT segment and MG3 domain of C3, as seen with C2a, Bb also contacts C3-MG8 via several polar residues uniquely present in its loop 2. The β-hairpin comprising the first two β-strands in the Bb-SP domain are longer than the corresponding region in C2a-SP and effectively packs onto Asn750_C3_–Asp752_C3_ (P2’–P4’ residues) by inducing the formation of a short strand (Fig. 4D, Fig. S5D). The tip of this β-hairpin also contacts C3-MG3. A hydrophobic pocket formed by Pro570_Bb_ and Ala663_Bb_–Tyr666_Bb_ accommodates Leu744_C3_ and Leu746_C3_ (Fig. 4E). Gly720_Bb_–Asp723_Bb_ at the beginning of loop 2 holds Arg748_C3_ in a similar manner as Gly700_C2a_–Asn703_C2a_, with the positive charge of Arg748_C3_ better neutralized by Asp723_Bb_. The Asp576_Bb_–His526_Bb_–Ser699(Ala)_Bb_ triad adopts nearly identical positions to Asp561_C2a_– His507_C2a_–Ser679(Ala)_C2a_, ready to attack the Arg748_C3_–Ser749_C3_ peptide bond.

## Stabilization of C3bBb by properdin

Properdin is the only known positive regulator of the complement pathway. Previous studies have demonstrated that properdin primarily interacts with the C345C domain of C3b through two critical loops: the TSR5-stirrup (or thumb) and TSR6-stirrup (or index finger) (*28, 29*). Focused refinement in this region allows us to pinpoint how these two loops insert into the C3b–Bb interface (Fig. 5A). Notably, Arg330_P_ in the TSR5-stirrup loop forms a salt bridge with both Asp1534_C3b_ and the main-chain oxygen of Leu1533_C3b_, effectively positioning itself between Leu1533_C3b_–Asp1534_C3b_ on one side and Pro1662_C3b_ on the other. This arrangement exerts pressure on Pro1662_C3b_ and facilitates the engagement of the terminal Asn1663_C3b_ with the metal ion (Ni^2+^ in this case) in the MIDAS motif of the Bb-vWA domain (Fig. 5B). Additionally, Arg329_P_ in the TSR5-stirrup forms a hydrogen bond with the main-chain oxygen of Leu349_Bb_, while stacking on Phe1659_C3b_ as previously described (Fig. 5C) (*29*). Ile340_P_ and Pro341_P_ engage Leu349_Bb_ through hydrophobic interactions. In the TSR6-stirrup, Ser419_P_ and Met420_P_ interact with Tyr317_Bb_ and Lys350_Bb_ via their main-chain oxygens. Collectively, these interactions from properdin serve to glue Bb onto C3b, thereby stabilizing the alternative pathway C3 convertase.

**Fig. 5:**
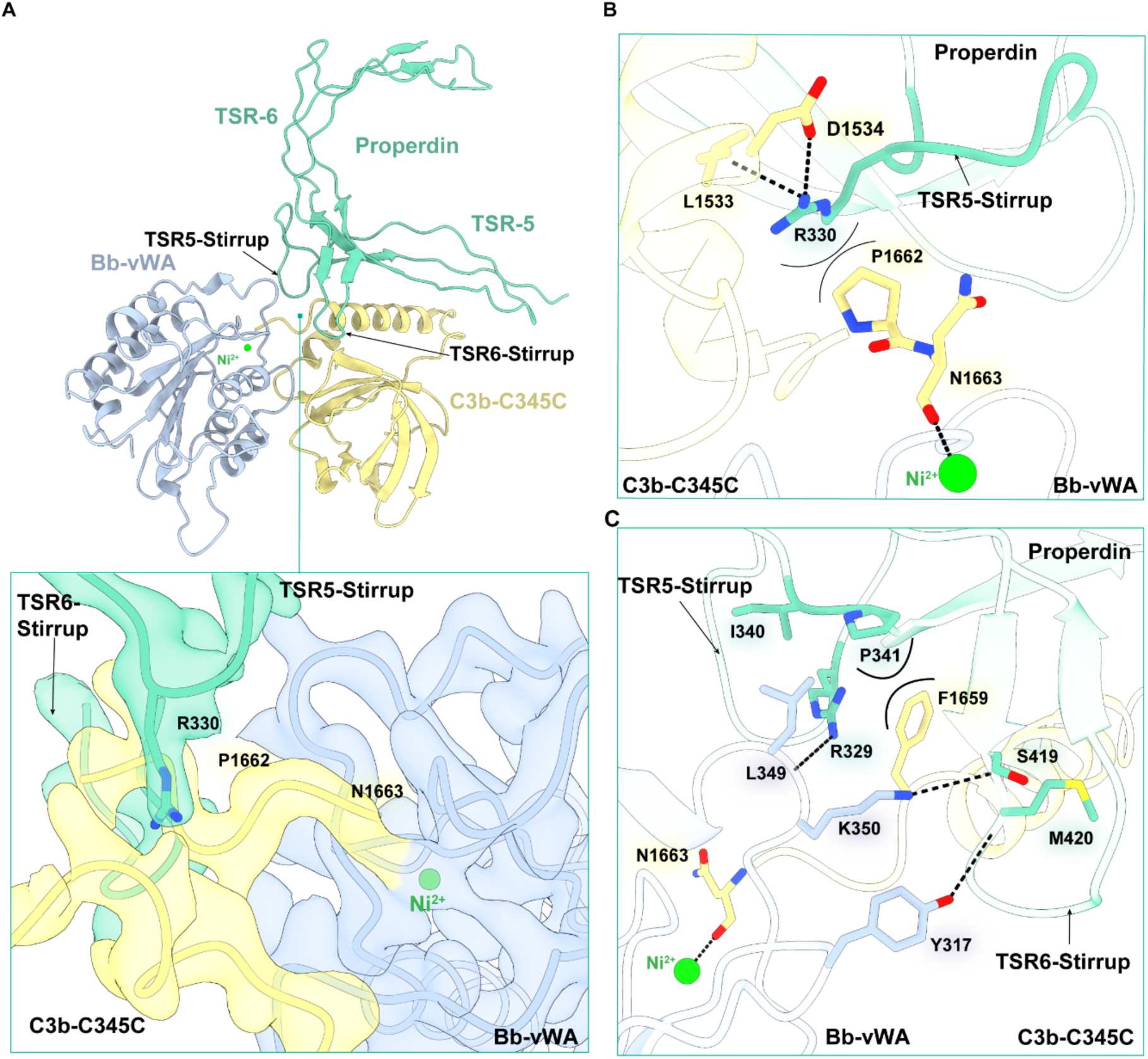
Stabilization of C3bBb by properdin. (A) The TSR5-stirrup and TSR6-stirrup loops of properdin promote the interaction between C3b and Bb. Bb-vWA, C3b-C345C, and properdin are shown in light blue, khaki and aquamarine, respectively. The density map of the structural interface is shown below. (B) Arg330_P_ facilitates the interaction between C3b and the metal ion in the MIDAS motif of Bb-vWA. (C) Additional interactions between properdin and C3bBb.

## Discussion

Targeting the complement C3 convertase represents a promising therapeutic strategy for various diseases, particularly those linked to complement dysregulation, such as certain kidney disorders. Pegcetacoplan, a member of the compstatin family of cyclic peptide C3 inhibitors, has been approved for treating paroxysmal nocturnal hemoglobinuria (PNH) and geographic atrophy (*30, 31*). Structural studies have shown that the compstatin molecules bind at the MG4– MG5 interface of C3 (*32, 33*). Consistent with previous analyses, compstatin molecules like Cp40, when engaged with C3, hinder the binding of C3 to either C4b2a or C3bBb, effectively blocking C3 activation across all complement pathways (Fig. 6A; Fig. S6). Notably, Cp40 can also bind to C3b, inhibiting the interaction between C3bBb and its C3 substrate, providing an additional layer of inhibitory effect. This dual mechanism makes compstatins ideally suited for broad-spectrum complement inhibition (Fig. 6A). Iptacopan, a factor B inhibitor, is another therapeutic agent used for treating PNH. Structural comparisons of the Bb-SP–Iptacopan complex (*34*) with our C3bBbP–C3 structure reveal that Iptacopan directly obstructs the entry of the α’-NT segment into the active site (Fig. 6B).

**Fig. 6:**
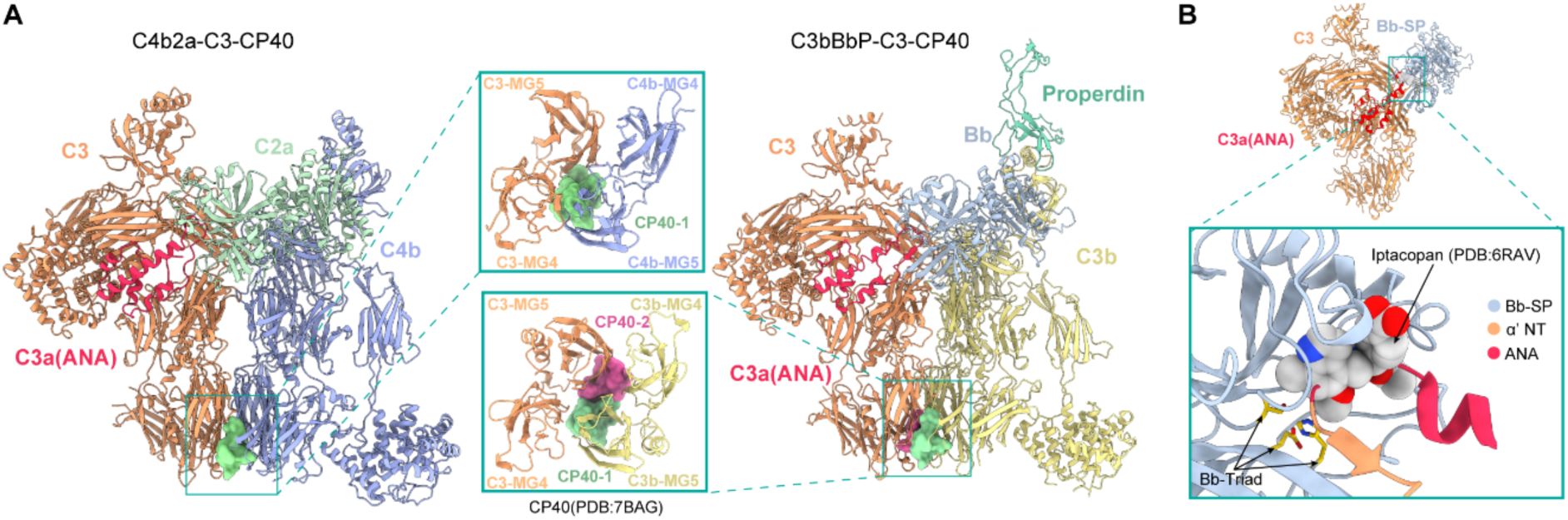
Insights into therapeutic drugs targeting the C3 convertase. **(A)** Mechanism of CP40 inhibition on the engagement of C3 with C4b2a and C3bBb. The C4b2a–C3 inhibition model is generated by superimposing C3b-Cp40 (PDB: 7BAG) onto the C3 structures of the C4b2a–C3 complex (left panel). The C3bBbP–C3 inhibition model is created by superimposing C3b-Cp40 onto both C3b and C3 (right panel). The binding of CP40 would create steric clashes that hinder C3 binding to either C4b2a or C3bBb. **(B)** Mechanism of Iptacopan inhibition on the engagement of C3 with C3bBbP. The inhibition model is generated by superimposing Bb-Iptacopan (PDB: 6RAV) onto the structure of Bb- C3a-α’-NT, which is separated from the C3bBbP–C3 complex. Iptacopan is represented in space-filling format.

Additionally, ongoing research is directed towards developing antibodies or nanobodies that specifically target C3 or the C3 convertases, examining these molecules in the context of our structural models to confirm their proposed mechanisms. For instance, both the S77 antibody and the hC3Nb1 nanobody bind to the C3b-MG7 domain (*35, 36*), disrupting the interaction between the C3bBb convertase and C3 (Fig. S6C). Furthermore, C4b-targeting nanobodies, such as NbE3, inhibit the C4b–C2a and C4b2a–C3 interactions, respectively (Fig. S6D) (*37*).

In summary, we have delineated the molecular mechanisms by which C3 is recognized by both C4b2a and C3bBb. Our results highlight distinct substrate-binding characteristics of the classical/lectin and alternative convertases and elucidate the unique role of properdin in the alternative pathway. These findings provide a fundamental understanding of these essential molecular machineries and will guide the design and optimization of future therapeutic molecules aimed at modulating complement activation.

## Acknowledgments

We are grateful to the Cryo-EM and the high-performance computing platforms of Peking University and Changping Laboratory for support with data collection and computation. We also thank the National Center for Protein Sciences at Peking University for assistance with the AKTA facility.

## Author contributions

C.J. and X.Y. made equal contributions to this work. Protein purification and complex assembling: C.J., X.Y.

Cryo-EM sample preparation, data acquisition, image processing, structure determination: X.Y., C.J.

Funding acquisition: J.X., Y.T., M.H.Z. Project administration: J.X., Y.T., M.H.Z. Supervision: J.X., Y.T., M.H.Z.

Writing – original draft: C.J., X.Y.

Writing – review & editing: J.X., Y.T., M.H.Z.

## Data and materials availability

Cryo-EM density maps have been deposited in the Electron Microscopy Data Bank with accession codes EMD-63893 (C4b2 in loading state), EMD-63892 (C4b2 in activation state), EMD-63894 (C4b2a–C3 complex) and EMD-63895 (C3bBbP–C3 complex). Structural coordinates have been deposited in the Protein Data Bank (PDB) with the accession codes 9U60, 9U5Z, 9U61 and 9U62. The other structural coordinates involved in this study are available from PDB (2ODP, 2WIN, 6S0B, 6EHG, 3G6J, 6RAV, 1ELV, 5JPM, 2XWB and 3HS0).

## Materials and Methods

### Cell line

#### HEK293F cells

HEK293F cells (Thermo) were cultured in FreeStyle 293 Expression Medium (Thermo) at 37°C with 5% CO₂ and 55% humidity using a humidified shaker for protein purification.

### Protein purification

#### Human C3 and C4 purification

Human C3 and C4 were purified from plasma provided by Peking university first hospital renal department as previously described (*19*).

For C3 purification, Na₂SO₄ powdered salt crystals (Sigma-Aldrich) were added to the plasma to a final concentration of 10% (w/v) to precipitate high-molecular-weight components. Following centrifugation and dialysis of the supernatant against the DEAE buffer, the protein solution was loaded onto a DEAE ion exchange column for the initial separation step. The C3- containing fraction was then dialyzed against Mono S column buffer and applied to a Mono S column to remove C3(H₂O). The eluted protein was concentrated using a 10,000-molecular-weight cutoff concentrator (Millipore) and further purified by size-exclusion chromatography using a Superdex 200 Increase column. The C3 fraction was collected, centrifuged, and finally stored in 25 mM HEPES-NaOH buffer (pH 7.4) with 100 mM NaCl.

For C4 purification, trisodium citrate was added to the plasma to a final concentration of 25 mM, followed by the slow addition of BaCl₂ solution to reach a concentration of 60 mM. The resulting precipitate was removed by centrifugation, and the supernatant was applied to a Q FF ion exchange column. The pooled eluent was treated with a 50% PEG 6000 solution to induce protein precipitation. After redissolving the protein, the solution was loaded onto a Q HP column. The fractions containing C4 were collected and further purified using a Mono Q column. Finally, C4 was subjected to gel filtration using a Superdex 200 Increase column for the final purification step and stored in 25 mM HEPES-NaOH buffer (pH 7.4) containing 150 mM NaCl.

#### Generation of C3b and C4b

C3b was generated by limited trypsin processing of C3 as previously described (*26*). Firstly, trypsin was added in a trypsin:C3 mass ratio 1:40 to cleave C3 at 37°C for 6 min. Subsequently, sample was transferred on ice immediately and soybean trypsin inhibitor was added to stop the reaction. C3b sample was subjected to gel filtration using a Superdex 200 Increase column for the final purification step and stored in 25 mM HEPES-NaOH buffer (pH 7.4) containing 100 mM NaCl. C4b was produced by adding active C1s (CompleTech., Cat. No. A104) to C4 solution in mass ratio 1:100, incubation first at 37 ℃ for 3-4 h and then overnight at 4 ℃.

#### Recombinant protein expression and purification

For recombinant expression of C2, Factor B, and Factor D in HEK293F cells, the genes encoding full-length C2 (UniProt P06681), Factor B (UniProt P00751) and Factor D (UniProt P00746) were cloned into the pcDNA3.1 vector with an 8×His-tag. For obtaining inactivated Factor B and C2 variants, site-directed mutagenesis was performed to introduce the S699A mutation in Factor B and the S679A mutation in C2.

HEK293F cells were cultured to a density of 1×10⁶ cells per milliliter and transfected with plasmids encoding the target proteins. For a 1 L cell culture, 1 mg of plasmid DNA was mixed with 2 mg of 40-kDa linear polyethyleneimine (PEI, Polysciences) in 50 mL of fresh medium and incubated for 20 minutes before transfection. The transfected cells were cultured for 4 days, after which the supernatant was collected by centrifugation. The medium was then exchanged with purification buffer (25 mM Tris-HCl, pH 7.4, and 150 mM NaCl) using a Hydrosart ultrafilter (Sartorius). Recombinant proteins were isolated by Ni-NTA affinity chromatography and eluted with purification buffer supplemented with 300 mM imidazole. The eluted proteins were further purified by size-exclusion chromatography using a Superdex 200 Increase column and stored in 25 mM HEPES-NaOH buffer (pH 7.4) with 150 mM NaCl.

A TEV site was introduced between residue P255 and V256 in properdin (Uniprot P27918). Subsequently, this properdin coding DNA was cloned into pcDNA3.1 vector with an 8×His-tag. After expression and purification from HEK293F cells, TEV protease was added in purified properdin in a TEV:protein mass ratio of 1:20 and incubated for 24 h at 4°C. The cleaved protein was further purified by size-exclusion chromatography using a Superdex 200 Increase column and stored in 25 mM HEPES-NaOH buffer (pH 7.4) with 150 mM NaCl.

#### Cryo-EM sample preparation and data collection

To prepare the classical pathway C3 convertase-substrate complex, C4b, C2(S679A) were mixed at a 1:1 molar ratio in binding buffer (25 mM HEPES-NaOH, pH 7.4, 100 mM NaCl, 2 mM NiCl_2_) and C1s was then added in a C2:C1s mass ratio 1:20. After incubation for 5 minutes at room temperature, the mixture was cooled to 4 °C and an equimolar amount of C3 was added. After a 30-minute incubation on ice, the solution was concentrated to a final protein concentration of 1 mg/mL using a 10,000-molecular-weight cutoff concentrator (Millipore).

To prepare the alternative pathway C3 convertase-substrate complex, C3b, CFB(S699A) and properdin-TEV were mixed at an equal molar ratio in binding buffer (25 mM HEPES-NaOH, pH 8.0, 100 mM NaCl, 2 mM NiCl_2_). Subsequently, Factor D was added in a Factor B: Factor D mass ratio 1:20. After incubation for 5 minutes at room temperature, the mixture was then cooled to 4 °C and an equimolar amount of C3 was added. After a 30-minute incubation on ice, the solution was concentrated to a final protein concentration of 0.7 mg/mL using a 10,000-molecular-weight cutoff concentrator (Millipore).

To prepare the C4b2b–C3 cryo-EM sample, 4 μL of the sample was deposited onto glow-discharged holey-carbon gold grids (Quantifoil Au 300 mesh R 0.6/1), which had been treated for 40 seconds using a Solarus Model 950 Plasma Cleaner (Gatan). The grids were blotted for 2.5 second with a blotting force of −1 at 4°C and 100% humidity, then plunged into liquid ethane cooled by liquid nitrogen. For the C3bBbP–C3 sample, 4 μL of the sample was deposited onto glow-discharged holey-carbon grids with continuous carbon film (Quantifoil Cu 300 mesh R 1.2/1.3 with 2nm C), which had been treated for 40 seconds using a Solarus Model 950 Plasma Cleaner (Gatan). The grids were blotted for 2 second with a blotting force of −1 at 4°C and 100% humidity, then plunged into liquid ethane cooled by liquid nitrogen.

All grids were initially screened using a 200 kV Talos Arctica microscope equipped with a Falcon 4 camera. Data collection was performed using EPU software (Thermo Scientific) on a 300 kV Titan Krios G4 microscope equipped with a Falcon 4 camera.

### Cryo-EM data processing

Data for determining the structures of C4b2b–C3, C4b2 (activation), C4b2 (loading) was from the same data set. A total of 14,681 movies were collected and processed by cryoSPARC (version 4.4.1) (*38*). Movie stacks were motion-corrected using the patch motion correction, and the CTF parameters were determined using the patch CTF estimation. Preliminary processed images were then manually screened to remove low-quality images through exposure curation. Subsequently, particle picking was performed using a blob picker, and templates were generated through subsequent 2D classification. The particles from template picking were subjected to multiple rounds of 2D classification to exclude inaccurate particles. Then the particles belonged to several kinds of 2D classification templates were selected and input to topaz train. The output model of topaz train was applied to selected more accurate particles and performed several rounds heterogeneous refinement to select the appropriate particles. The C4b2b–C3 particles resulting from the heterogeneous refinement were then utilized in homogeneous refinement and local refinement for the generation of the final 3D reconstruction. The C4b2 (loading state) and C4b2 (activation state) particles resulting from the heterogeneous refinement were then carried out several 3D classifications. To obtain a higher-quality local density map, mask-based local refinement was further performed using cryoSPARC. The local resolution map was produced with the local resolution estimation program in cryoSPARC. Data of C4b2b–C3, C4b2 (loading state) and C4b2 (activation state) were processed in a same way. Finally, 109,455 C4b2b–C3 particles, 109,606 C4b2 (loading state) particles, 36,092 C4b2 (activation state) particles were used to reconstruction and yielded a map of 3.09 Å, 2.91 Å, 3.06 Å resolution, respectively.

For the C3bBbP–C3 complex, a total of 8,649 movies were collected and processed by cryoSPARC (version 4.4.1). Movie stacks were processed and particle were picked in a similar protocol. The particles from topaz extract were performed several rounds heterogeneous refinement to select the appropriate particles. The C3bBbP–C3 particles resulting from the heterogeneous refinement were then utilized in homogeneous refinement and local refinement for the generation of the final 3D reconstruction. Mask-based focus refinement was further performed using cryoSPARC to obtain the higher-quality local density maps for model building. Finally, 389,342 particles, 93,849 particles, 194,378 particles were used to the global map, C3b-TED-focused local map, and P-focused local map reconstruction and yielded a resolution of 2.57 Å, 2.96 Å and 2.73 Å, respectively.

### Model building and structure refinement

Initial models of C4b, C3 and C2 were generated using Alphafold2 (*39*). Models of C2a, C3b, Bb and properdin was obtained from previous structure (PDB ID: 2ODP, 2WIN, 6S0B) (*18, 29, 40*). These models were docked into the cryo-EM density map using UCSF Chimera (*41*). Further structure model building was performed using Coot (*42*) and refined using the real-space refinement in Phenix (*43*). Figures were prepared with UCSF ChimeraX (*44*).

---

**Fig. S1:**
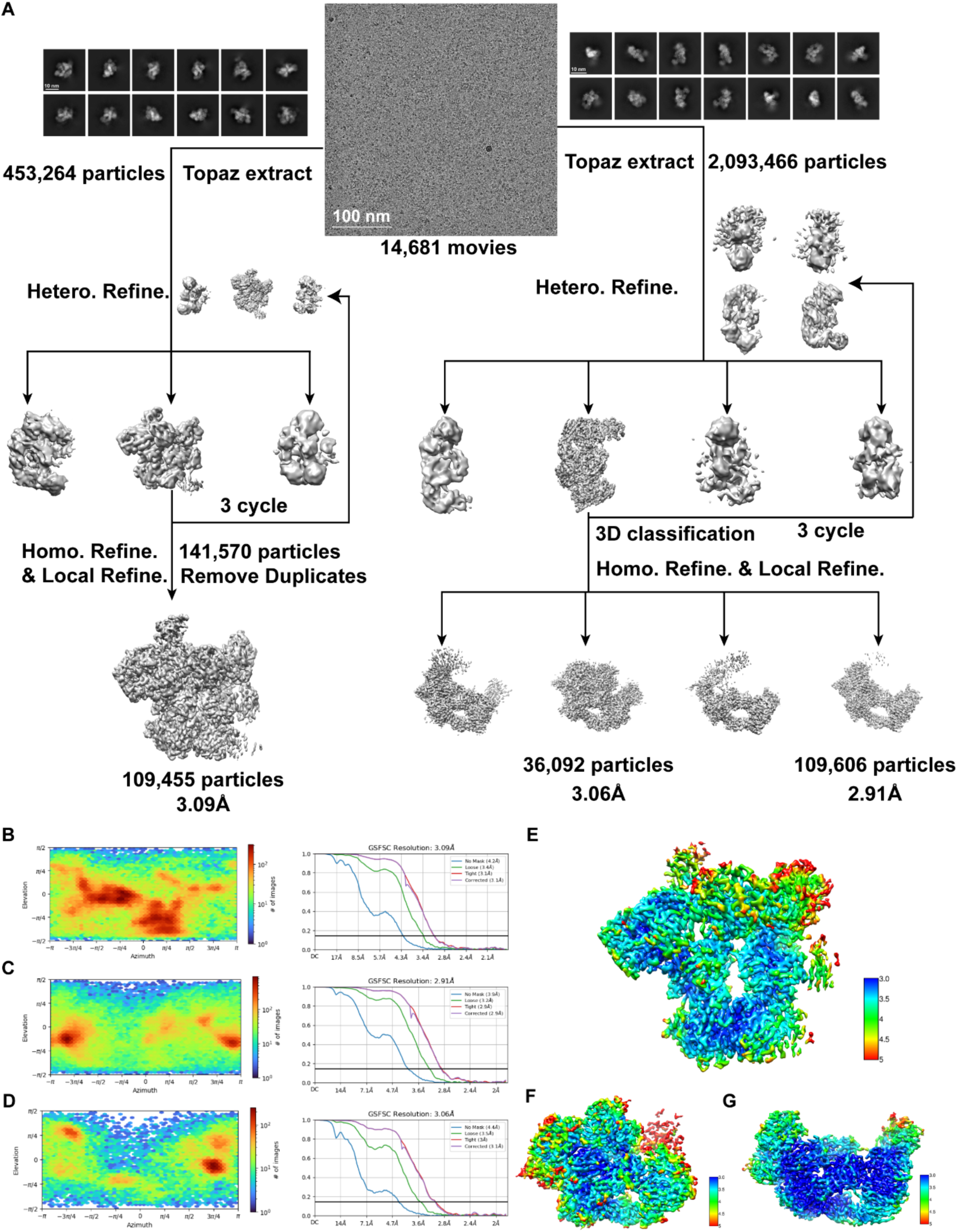
Cryo-EM 3D reconstruction of the C4b2a–C3 complex, C4b2(loading) and C4b2(activation). **(A)** Flowchart of cryo-EM data processing for C4b2a–C3, C4b2 (loading) and C4b2(activation). The density maps of C4b2a–C3 and C4b2 were generated from the same data set. **(B)-(D)** Angular particle distribution heat map and Gold-standard Fourier shell correlation (GSFSC) curves of C4b2a–C3, C4b2 (loading) and C4b2(activation), respectively. **(E)-(G)** Resolution estimations for the final maps of C4b2a–C3, C4b2 (activation) and C4b2 (loading), respectively.

**Fig. S2:**
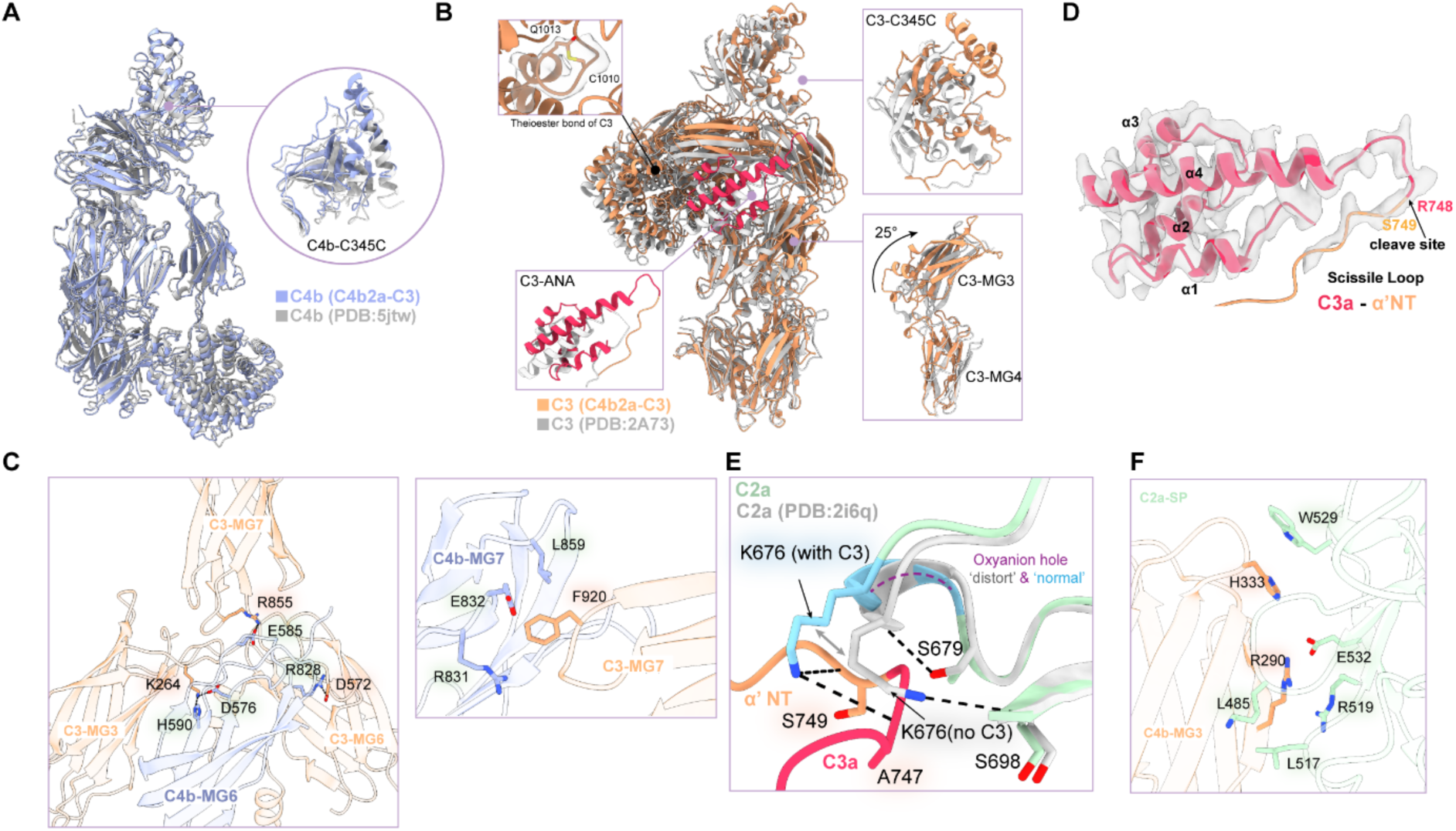
Structure analysis of C4b2a–C3 complex. **(A)** C4b structure in the C4b2a–C3 complex compared to C4b alone (PDB: 5jtw). **(B)** C3 structure in the C4b2a–C3 complex compared to C3 alone (PDB: 2A73). **(C)** C4b-MG6 and C4-MG7 interact with C3-MG3, C3-MG6, and C3-MG7. **(D)** Density map of C3-ANA and α’-NT. Black arrow indicates the cleavage site between R748 and S749. **(E)** A functional oxyanion hole is established in C2a-SP upon C3 binding, when compared to the C2a alone structure (PDB: 2i6q). **(F)** Interaction between C2a-SP and C4b-MG3.

**Fig. S3:**
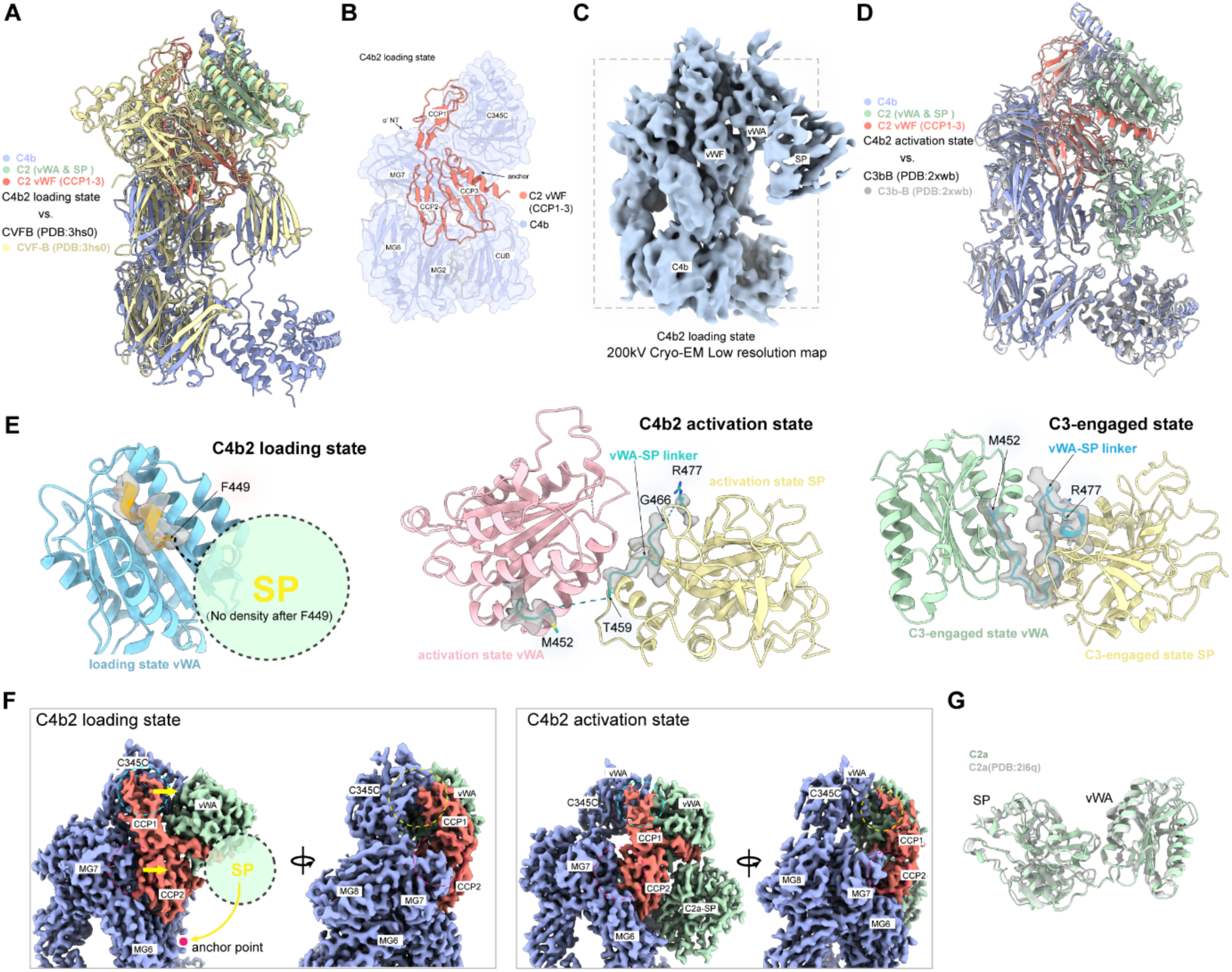
Activation of classical pathway C3 convertase. **(A)** Structural comparison of CVF-B (PDB: 3hs0) and the loading state of C4b2. CVF-B is shown in yellow, and C4b2 is shown in purple, green and salmon. **(B)** Interaction between the vWF domain of C2 and C4b in the loading state. **(C)** A low resolution density map of the C4b2 loading state reveals the general location of the C2a-SP. **(D)** Structural comparison of C3bB (PDB: 3xwb) and the activation state of C4b2. C3bB is shown in grey, and the C4b2 activation state structure is shown in purple, green and salmon. **(E)** The linker between C2a-vWA and C2a-SP domain undergo conformational changes from the loading state to activation state, and subsequently to C3-bound state. **(F)** The docking of C2-SP onto C4b results in reduced interaction between C2a-vWF and C4b- C345C, facilitating the detachment of C2a-vWF. **(G)** Structure comparison of C2a in the C3-bound state and the C2a alone structure (PDB: 2i6q).

**Fig. S4:**
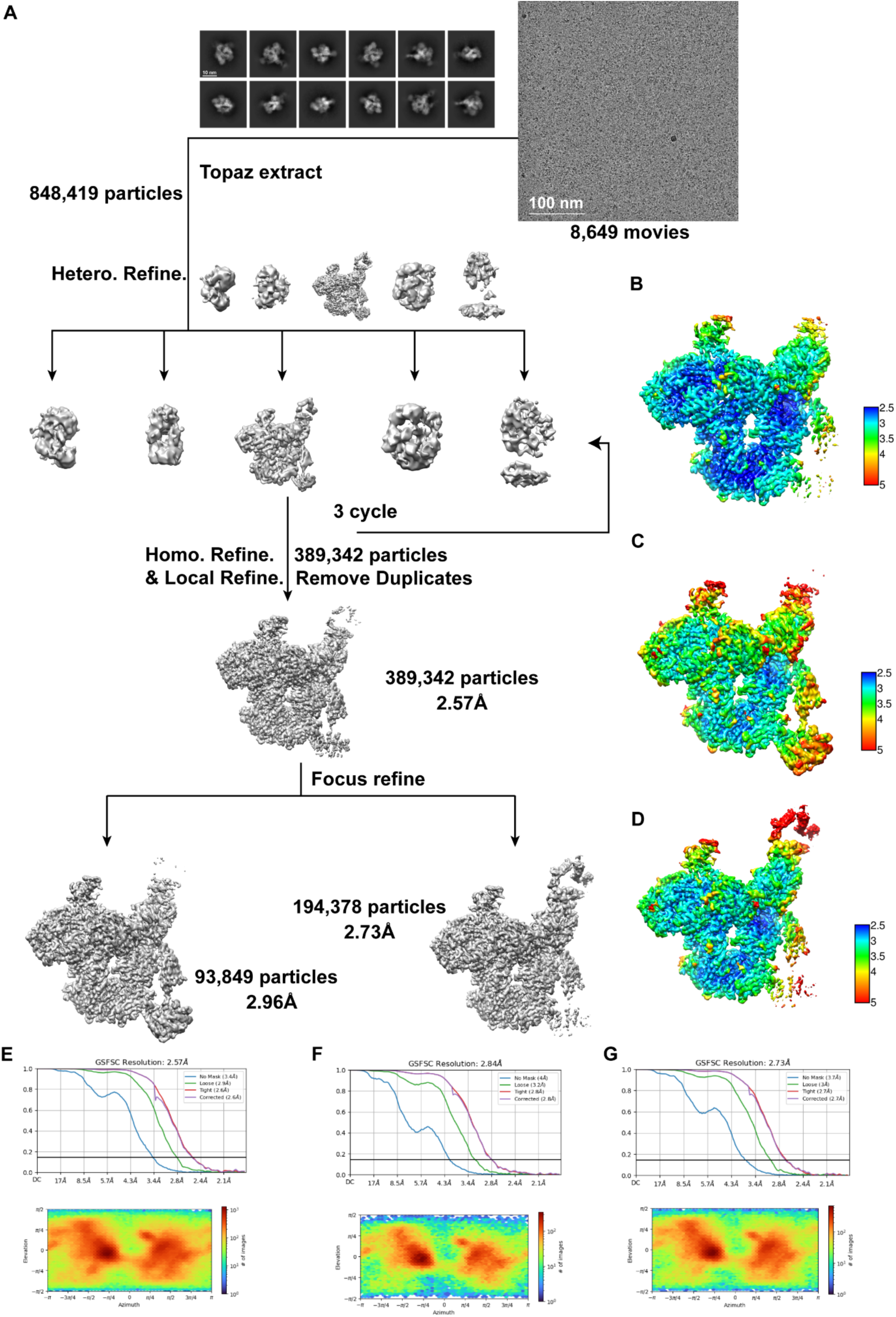
Cryo-EM 3D reconstruction of the C3bBbP–C3 complex. **(A)** Flowchart of cryo-EM data processing for the C3bBbP–C3 complex. **(B)-(D)** Resolution estimations for the final maps of the C3bBbP–C3 complex and the focused map of C3bBbP–C3 complex, respectively. **(E)-(G)** Angular particle distribution heat map and Gold-standard Fourier shell correlation (GSFSC) curves of the C3bBbP–C3 complex and the focused map of C3bBbP–C3 complex, respectively.

**Fig. S5:**
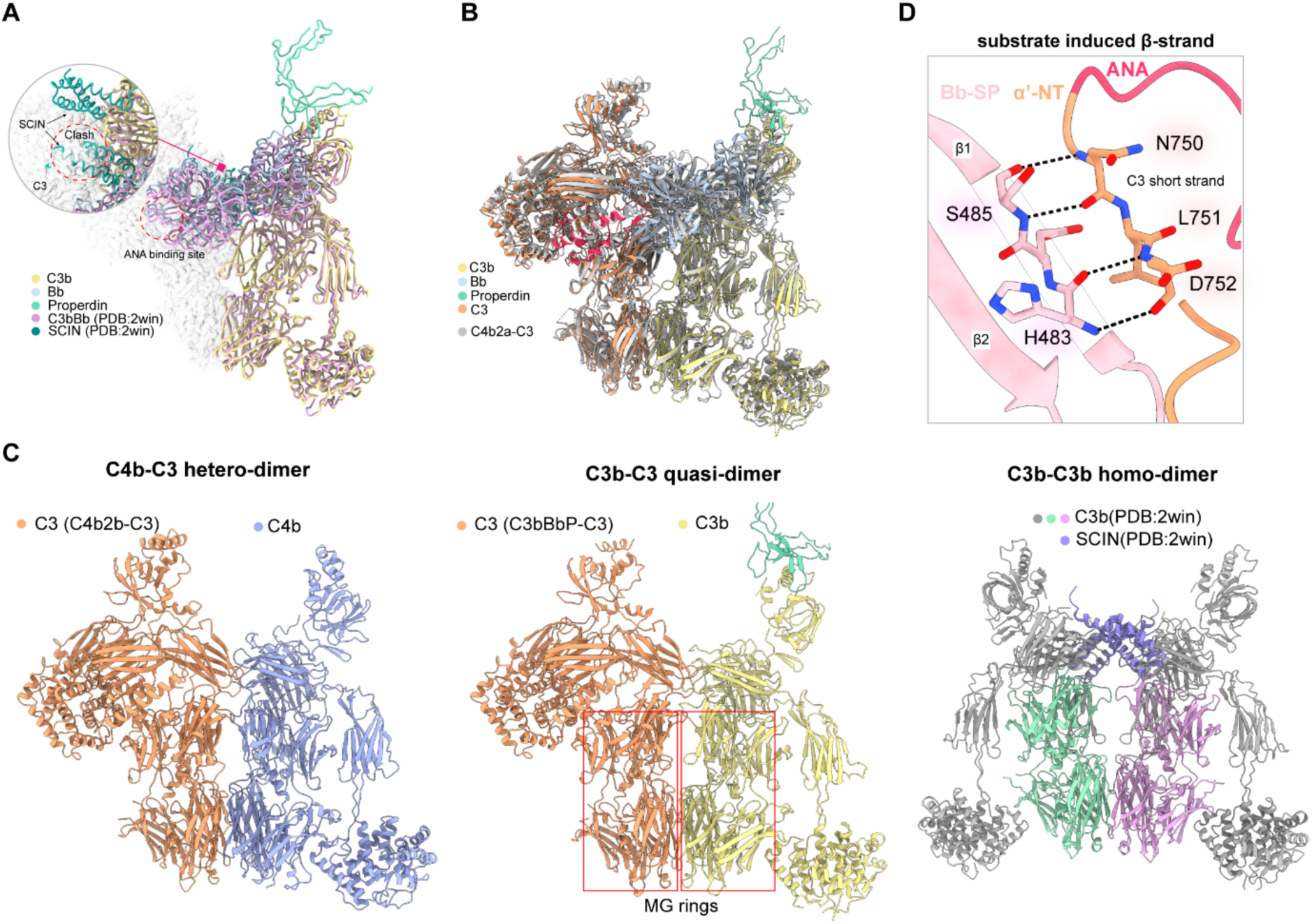
Cryo-EM structure analysis of the C3bBbP–C3 complex. **(A)** Structural superimposition of the C3bBbP–C3 and C3bBb–SCIN (PDB: 2win). C3bBb– SCIN is shown in pink. C3 in the C3bBbP–C3 complex is shown using a surface representation. The presence of SCIN would prevent the binding of C3 to C3bBb due to steric hindrance. **(B)** Structural superimposition of C3bBbP–C3 and C4b2a–C3. **(C)** Structural comparison between the C4b-C3 heterodimer, C3b-C3 quasi-dimer and C3b- C3b homodimer. Red boxes indicate MG-rings within C3 and C3b. **(D)** Interactions between the Bb-SP hairpin structure and C3 α’-NT.

**Fig. S6:**
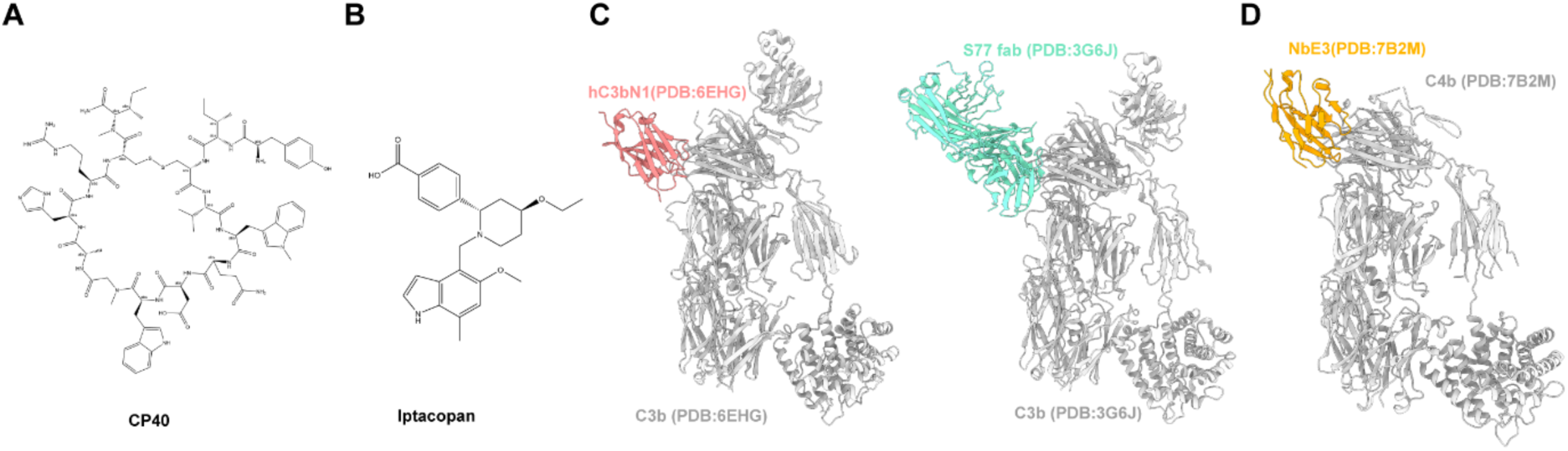
C3 inhibitory molecules. **(A)** Chemical structure of the cyclic peptide CP40. **(B)** Chemical structure of the Iptacopan (Fabhalta). **(C)** Structure of hC3bN1-C3b and S77-C3b. hC3bN1 is a C3b-MG7 binding nanobody, whereas S77 is a C3b-MG7 binding antibody. Both hC3bN1 and S77 would inhibit C3bBb by obstructing the binding of the substrate C3 molecule. **(D)** Structure of NbE3-C4b. NbE3 would block the binding C3 to C4b and thereby inhibit the C4b2a convertase.

**Table S1.** Cryo-EM data collection, refinement and validation statistics.

|  | C4b2_loading | C4b2_activation | C4b2a_C3 | C3bBbP_C3 |
| --- | --- | --- | --- | --- |
| **Data collection and processing** |  |  |  |  |
| Magnification |  | 130,000 |  |  |
| Voltage (kV) |  | 300 |  |  |
| Electron exposure (e-/Å<sup>2</sup>) |  | 60 |  |  |
| Defocus range (μm) |  | -1.1 to -1.5 |  |  |
| Pixel size (Å) |  | 0.95 |  |  |
| Symmetry imposed |  | C1 |  |  |
| Initial particle images (no.) | 2,093,466 | 2,093,466 | 453,264 | 848,419 |
| Final particle images (no.) | 109,606 | 36,0922 | 109,455 | 389,342 |
| Map resolution (Å) |  |  |  |  |
| FSC threshold: 0.143 | 2.91 | 3.06 | 3.09 | 2.57 |
| **Refinement** |  |  |  |  |
| Initial model used (PDB code) | Alphafold2 | Alphafold2 | Alphafold2<br>2ODP | Alphafold2<br>2WIN, 6S0B |
| Model resolution (Å) |  |  |  |  |
| FSC threshold: 0.143 | 2.91 | 3.06 | 3.09 | 2.57 |
| Map sharpening *B* factor (Å<sup>2</sup>) | 62.0 | 44.0 | 59.6 | 83.9 |
| Model composition |  |  |  |  |
| Non-hydrogen atoms | 14,719 | 16,447 | 29,000 | 29,466 |
| Protein residues | 1,899 | 2,116 | 3,674 | 3,709 |
| Ligands | 1 | 1 | 1 | 1 |
| *B* factors (Å<sup>2</sup>) |  |  |  |  |
| Protein | 81.19 | 122.56 | 92.51 | 458.80 |
| Ligand | 106.13 | 130.72 | 100.96 | 127.37 |
| R.m.s. deviations |  |  |  |  |
| Bond lengths (Å) | 0.004 | 0.007 | 0.005 | 0.004 |
| Bond angles (°) | 0.813 | 1.141 | 0.837 | 0.829 |
| Validation |  |  |  |  |
| MolProbity score | 1.62 | 1.76 | 1.41 | 1.69 |
| Clashscore | 2.87 | 4.74 | 3.15 | 3.25 |
| Poor rotamers (%) | 0.56 | 0.28 | 0.41 | 3.30 |
| Ramachandran plot |  |  |  |  |
| Favored (%) | 95.67 | 91.05 | 95.65 | 96.92 |
| Allowed (%) | 4.33 | 8.95 | 4.33 | 3.08 |
| Disallowed (%) | 0.00 | 0.00 | 0.03 | 0.00 |

